# Mitochondrial Sodium/Calcium Exchanger (NCLX) Regulates Basal and Starvation-Induced Autophagy Through Calcium Signaling

**DOI:** 10.1101/2023.03.17.533187

**Authors:** Vitor de Miranda Ramos, Julian D.C. Serna, Eloisa A. Vilas-Boas, João Victor Cabral-Costa, Fernanda M. da Cunha, Tetsushi Kataura, Viktor I. Korolchuk, Alicia J. Kowaltowski

## Abstract

Mitochondria shape intracellular Ca^2+^ signaling through the concerted activity of Ca^2+^ uptake via mitochondrial calcium uniporter, and efflux from by Na^+^/Ca^2+^ exchangers (NCLX). Here, we describe a novel relationship between NCLX, intracellular Ca^2+^, and autophagic activity. Conditions that stimulate autophagy *in vivo* and *in vitro,* such as caloric restriction and nutrient deprivation, upregulate NCLX expression in hepatic tissue and cells. Conversely, knockdown of NCLX impairs basal and starvation-induced autophagy. Similarly, acute inhibition of NCLX activity by CGP 37157 affects bulk and endoplasmic reticulum autophagy (ER-phagy) without significant impacts on mitophagy. Mechanistically, CGP 37157 inhibited the formation of FIP200 puncta and downstream autophagosome biogenesis. Inhibition of NCLX caused decreased cytosolic Ca^2+^ levels, and intracellular Ca^2+^ chelation similarly suppressed autophagy. Furthermore, chelation did not exhibit an additive effect on NCLX inhibition of autophagy, demonstrating that mitochondrial Ca^2+^ efflux regulates autophagy through the modulation of Ca^2+^ signaling. Collectively, our results show that the mitochondrial Ca^2+^ extrusion pathway through NCLX is an important regulatory node linking nutrient restriction and autophagy regulation.

## Introduction

Mitochondria are highly dynamic organelles that can adapt their functions to changes in the cellular environment (Liesa and Shirihai, 2013; Giacomello et al., 2020). In return, they provide signals that regulate cytoplasmic processes, leading to coordinated responses (da Cunha et al., 2015). Among these, calcium (Ca^2+^) signaling is known to regulate and be regulated by mitochondrial activity (Rizzuto et al., 2012). Indeed, the capacity to take up and release Ca^2+^ ions makes mitochondria able to directly participate in a myriad of cellular Ca^2+^-sensitive regulation pathways (Rizzuto et al., 2012; Giorgi et al., 2018).

Mitochondrial Ca^2+^ uptake is mediated by the mitochondrial Ca^2+^ uniporter (MCU) complex (De Stefani et al., 2011), while Ca^2+^ efflux can occur through two specific transporters: a mitochondrial H^+^/Ca^2+^ exchanger, with a disputed molecular identity (Giorgi et al., 2018; Patron et al., 2022); and the mitochondrial Na^+^/Li^+^/Ca^2+^ exchanger (NCLX), which releases matrix Ca^2+^ in exchange for Na^+^ ions. With the identification of the NCLX gene in 2010 (Palty et al., 2010), several studies uncovered important physiological functions of this transporter, including regulation of insulin secretion, cardiomyocyte automaticity, and lactate and glutamate secretion in astrocytes, all promoted by shaping Ca^2+^ signals (Nita et al., 2012; Takeuchi et al., 2013; Parnis et al., 2013; Cabral-Costa et al., 2023). In pathology, decreased NCLX expression was observed in Alzheimer’s disease, colorectal cancer, and heart failure patient tissues, while the rescue of its activity led to improved phenotypes (Jadiya et al., 2019; Pathak et al., 2020; Garbincius et al., 2022). These studies suggest that impaired NCLX activity is central in the regulation of pathways with extensive therapeutic potential.

Autophagy, a highly conserved intracellular pathway leading to lysosomal degradation of cellular components (Mizushima and Komatsu, 2011), is widely recognized as a pivotal process involved in maintaining tissue homeostasis. In macroautophagy (which we will refer to as autophagy from now on), these components are sequestered by autophagosomes, which in turn fuse with lysosomes to promote the digestion of cargo such as organelles or proteins (Mizushima and Komatsu, 2011). Due to its ability to remove unwanted elements and promote specific or general nutrient availability as an output, autophagy is necessary for quality control, tissue renovation, and metabolic regulation (Mizushima and Komatsu, 2011; Lahiri et al., 2019; Wilson et al., 2023).

Ca^2+^ ions are known second messengers in metabolic signaling and display an important role in autophagy regulation (Berridge and Lipp, 2000; Filippi-Chiela et al., 2016; Zhang et al., 2022). However, the genesis and consequences of Ca^2+^ signaling within this regulatory mechanism are very context-dependent and not yet fully understood (Decuypere et al., 2011a; Bootman et al., 2018). This is intrinsically related to the fact that Ca^2+^ signals are shaped at subcellular levels by ion transporters in different organelles. In fact, endeavors to study the role of specific Ca^2+^ transporters located in the endoplasmic reticulum (ER) or lysosome, rather than overall cellular Ca^2+^ levels, have strongly contributed toward untangling particular aspects of autophagic regulation (Medina et al., 2015; Decuypere et al., 2011b). Considering the presence of mitochondria in close contact with these organelles and the occurrence of Ca^2+^ transfer between them (Rizzuto et al., 1998; Raffaello et al., 2016; Peng et al., 2020), it is likely that mitochondrial Ca^2+^ transport participates in autophagy regulation. In fact, the transfer of Ca^2+^ from inositol trisphosphate receptor (IP_3_R) in the ER to mitochondria through the MCU was shown to suppress basal autophagy in an AMP-activated protein kinase (AMPK)-dependent manner (Cárdenas et al., 2010). On the other hand, the question of how mitochondrial Ca^2+^ efflux through NCLX can affect autophagy has not yet been addressed.

Thus, considering the emerging relevance of NCLX in a myriad of physiological and pathological processes in which autophagy is similarly important, and considering the importance of Ca^2+^ transporters in autophagic regulation, this work focused on investigating the interplay between NCLX activity and autophagy.

## Results and Discussion

### NCLX expression is increased by nutrient restriction

Autophagy is mainly activated in response to decreased nutrient availability (Lahiri et al., 2019). Interestingly, works published by our group have shown that mitochondrial Ca^2+^ transport is strongly regulated by nutritional interventions such as caloric restriction, which is also well known to promote autophagy (Menezes-Filho et al., 2017; Amigo et al., 2017; Serna et al., 2022). As a starting point to understand the interplay between mitochondrial Ca^2+^ transport and autophagy, we investigated whether NCLX levels are sensitive to calorie restriction. We consistently observed that mitochondria isolated from the livers of mice that underwent 4 months of caloric restriction contained higher levels of NCLX protein than animals on an *ad libitum* diet (**Figure 1A**). To test if this effect could be reproduced in an *in vitro* model of autophagy activation, we submitted AML12 cells, a non-tumoral cell line derived from murine hepatocytes (Wu et al., 1994), to serum and amino acid starvation using Hanks’ balanced salt solution (HBSS). Starvation resulted in a time-dependent increase in NCLX mRNA (**Figure 1B**). On the other hand, MCU mRNA levels were not changed under the same conditions (Fig S1), demonstrating that this effect is specific to NCLX, rather than a more global change in mitochondrial Ca^2+^ transport machinery. Overall, our results show that the specific expression of NCLX is highly sensitive to nutrient availability in hepatic cells, suggesting a physiological role in nutrient sensing.

**Figure 1.**
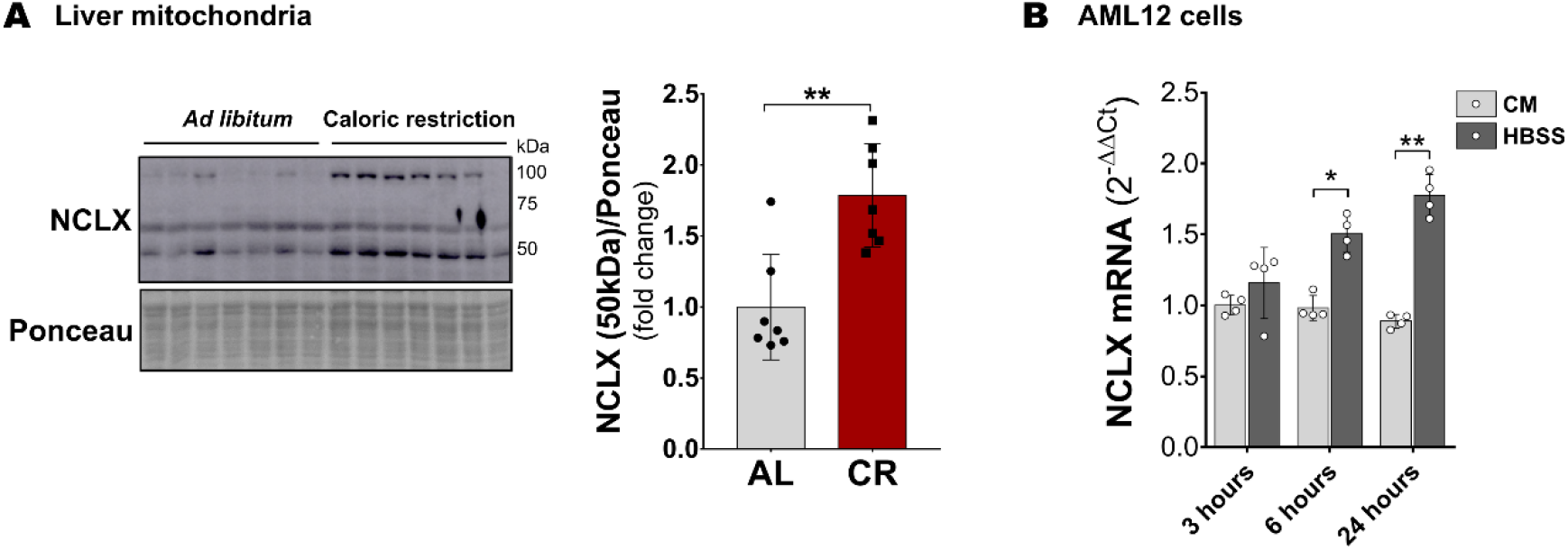
NCLX expression is increased by nutrient restriction. (**A**) Western blot analysis of NCLX protein levels in mitochondria isolated from the livers of Swiss mice after 4 months of *ad libitum* (AL) or caloric restriction (CR) diets (n = 7 per group). Data were analyzed by two-tailed unpaired t test. (**B**) RT-qPCR for NCLX mRNA in AML12 cells incubated with either complete media (CM) or HBSS for 3, 6, and 24 hours (n = 4). Data were analyzed by two-way RM ANOVA followed by Bonferroni’s multiple comparisons test. Bar graphs show means ± SD. Dots in each group represent biological replicates. * = p < 0.05, ** = p < 0.01, *** = p < 0.001, ns = not significant.

### Decreased NCLX expression causes impaired autophagic activity

Since we found NCLX expression to be increased during nutrient restriction, we tested whether reducing NCLX levels could affect starvation-induced autophagy by promoting NCLX genetic knockdown (KD) in hepatic AML12 cells transfected with NCLX-targeting siRNA (siNCLX). NCLX mRNA levels were significantly reduced 48 and 72 hours after transfection when compared to cells transfected with negative control siRNA (siNC) (**Figure 2A**). Consequently, NCLX protein levels were also decreased by siRNA-mediated KD (**Figure 2B**). As a readout of autophagic activity, we initially measured the levels of LC3 II and LC3 I proteins. As expected, siNC cells incubated with HBSS for 1 hour showed increased LC3 II and reduced LC3 I compared to cells incubated with complete media (CM), indicating autophagy activation (**Figure 2C**). In siNCLX cells, LC3 II levels were not upregulated by HBSS treatment (**Figure 2C**), which reflects an inhibition of autophagy. Curiously, LC3 I levels were lower in siNCLX cells even in the absence of starvation (**Figure 2C**), thereby suggesting a modulation of total LC3 by NCLX. Besides autophagic activity, LC3 levels may be affected by either proteasome degradation (Gao et al., 2010) or gene expression. We did not observe alterations in proteasome activity by NCLX KD (Fig S2A), while LC3A and LC3B mRNA were reduced compared to siNC (Fig S2B).

**Figure 2.**
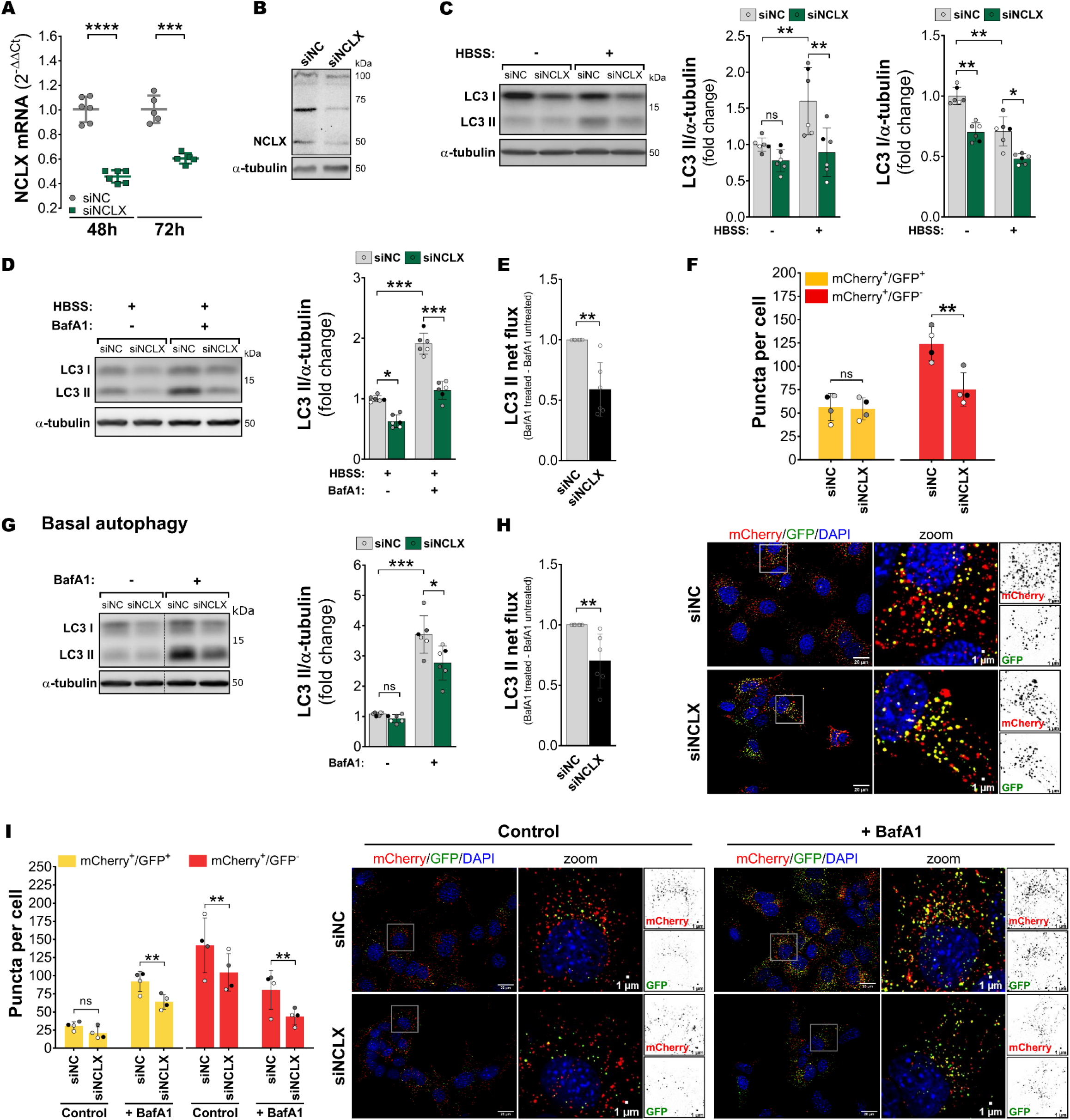
Decreased NCLX expression causes impaired autophagic activity. AML12 cells transfected with NCLX (siNCLX) or negative control (siNC) siRNAs were analyzed by: (**A**) RT-qPCR for NCLX mRNA, 48 (n = 6) and 72 (n = 5) hours after transfection. Data were analyzed by two-tailed paired t test. (**B**) Western blot of NCLX protein levels 72 hours after transfection. (**C**) Western blots for LC3 II and LC3 I after 1 hour incubation with either complete media (CM) or HBSS (n = 6). (**D**) Western blot of LC3 II after 1 hour incubation with HBSS in the absence or presence of 100 nM Bafilomycin A1 (BafA1) during the last 30 min of incubation (n = 6). Data were analyzed by two-way RM ANOVA followed by Tukey’s multiple comparisons test. (**E**) LC3 II net flux in siNC and siNCLX cells incubated with HBSS. Data were analyzed by two-tailed unpaired t test. (**F**) AML12 stably expressing the mCherry-EGFP-LC3B probe, showing representative images and quantification of autophagosomes (mCherry^+^/GFP^+^; yellow puncta) and autolysosomes (mCherry^+^/GFP^-^; red puncta) per cell after 1 hour incubation with HBSS (n = 4). Data were analyzed by two-tailed unpaired t test. (**G**) Western blot for LC3 II in CM conditions and in the absence or presence of 100 nM Bafilomycin A1 (+BafA1) for 1 hour (n = 6). Data were analyzed by two-way RM ANOVA followed by Tukey’s multiple comparisons test. (**H**) LC3 II net flux in siNC and siNCLX cells incubated with complete media. (**I**) AML12 stably expressing the mCherry-EGFP- LC3B probe, showing representative images and quantification of autophagosomes (mCherry^+^/GFP^+^; yellow puncta) and autolysosomes (mCherry^+^/GFP^-^; red puncta) per cell (n = 4). Data were analyzed by two-way RM ANOVA followed by Tukey’s multiple comparisons test. Data are presented in columns with means ± SD. Dots in each group represent biological replicates, and paired biological replicates are represented by equally colored symbols. * = p < 0.05, ** = p < 0.01, *** = p < 0.001, ns = not significant.

To confirm autophagy inhibition by NCLX KD, we repeated the experiment in the presence of the lysosomal inhibitor Bafilomycin A1 (BafA1). Consistently, cells with reduced NCLX expression showed significantly less LC3 II accumulation than siNC cells in the presence of BafA1 (**Figure 2D**). Accordingly, LC3 II net flux (BafA1-treated minus BafA1-untreated samples) was reduced in siNCLX cells (**Figure 2E**).

As an additional strategy, we used an AML12 cell line stably expressing mCherry-GFP- tagged LC3B, which allows the visualization of autophagosome (mCherry^+^/GFP^+^ puncta) and autolysosome (mCherry^+^/GFP^-^ puncta) formation using fluorescence microscopy (ŃDiaye et at., 2009; Yoshii and Mizushima, 2017). After 1 hour incubation in HBSS, siNCLX cells showed similar levels of autophagosomes to controls, but reduced autolysosome puncta (**Figure 2F**), hence supporting the finding that autophagic activity is compromised in these cells. Interestingly, since LC3 II levels were lower in siNCLX cells (**Figure 2C**), we expected autophagosome puncta to be similarly reduced, but did not see this in these cells, likely due to outcome interference by the LC3 probe system used. Nonetheless, our results with mCherry-GFP-tagged LC3B cells clearly demonstrate that reduced NCLX expression compromises autophagic activity during starvation.

Since reduced expression of NCLX led to changes in LC3 levels, we hypothesized that basal autophagy, in the presence of nutrients and serum, could be affected. Indeed, although steady-state (without BafA1) levels of LC3 II were unchanged, its accumulation in the presence of BafA1 was lower in siNCLX cells (**Figure 2G**), which also resulted in decreased net flux (**Figure 2H**). Additionally, siNCLX cells presented reduced autolysosome puncta, either under control conditions or with BafA1; autophagosome puncta were reduced after BafA1 treatment (**Figure 2I**).

Thus far, our results demonstrated that autophagic stimulation upregulates NCLX expression, while cells with reduced expression of this mitochondrial transporter have decreased autophagic activity and are less sensitive to starvation stimulus. Collectively, these findings indicate a direct relationship between NCLX levels and autophagic activity in hepatic cells.

### Acute NCLX inhibition impairs starvation-induced bulk and selective autophagy

Our data shown up to now indicate that chronic inhibition of NCLX affects autophagic machinery. However, the time frame required for silencing studies means the effect of NCLX KD on autophagy could be indirect. To address the hypothesis that Ca^2+^ efflux from the mitochondria could directly participate in the regulation of autophagy activation, we used the strategy of acute pharmacological inhibition of NCLX activity by CGP 37157 (CGP). To validate that the drug was effective under our conditions, we measured Ca^2+^ levels within the mitochondrial matrix using the genetically-encoded probe mito-GEM-GECO1 (Zhao et al., 2011). Cells incubated with HBSS in the presence of DMSO or 10 µM CGP were imaged upon the addition of 100 µM extracellular ATP to stimulate ER Ca^2+^ release (**Figure 3A**). ATP elicited mitochondrial Ca^2+^ uptake, resulting in a rapid increase in intramitochondrial Ca^2+^ levels followed by a slower decay towards basal levels (**Figure 3A, light grey line/column**), corresponding to the mitochondrial Ca^2+^ efflux rate. Ionomycin, a Ca^2+^ ionophore, was added at the end of all traces as an internal control. Importantly, cells in the presence of CGP (**Figure 3A, blue line/column**) had equal mitochondrial Ca^2+^ increase upon ATP stimulation (indicating no effect on ion uptake), but did not present a measurable efflux rate after the Ca^2+^ peak, thereby confirming specific NCLX inhibition.

**Figure 3.**
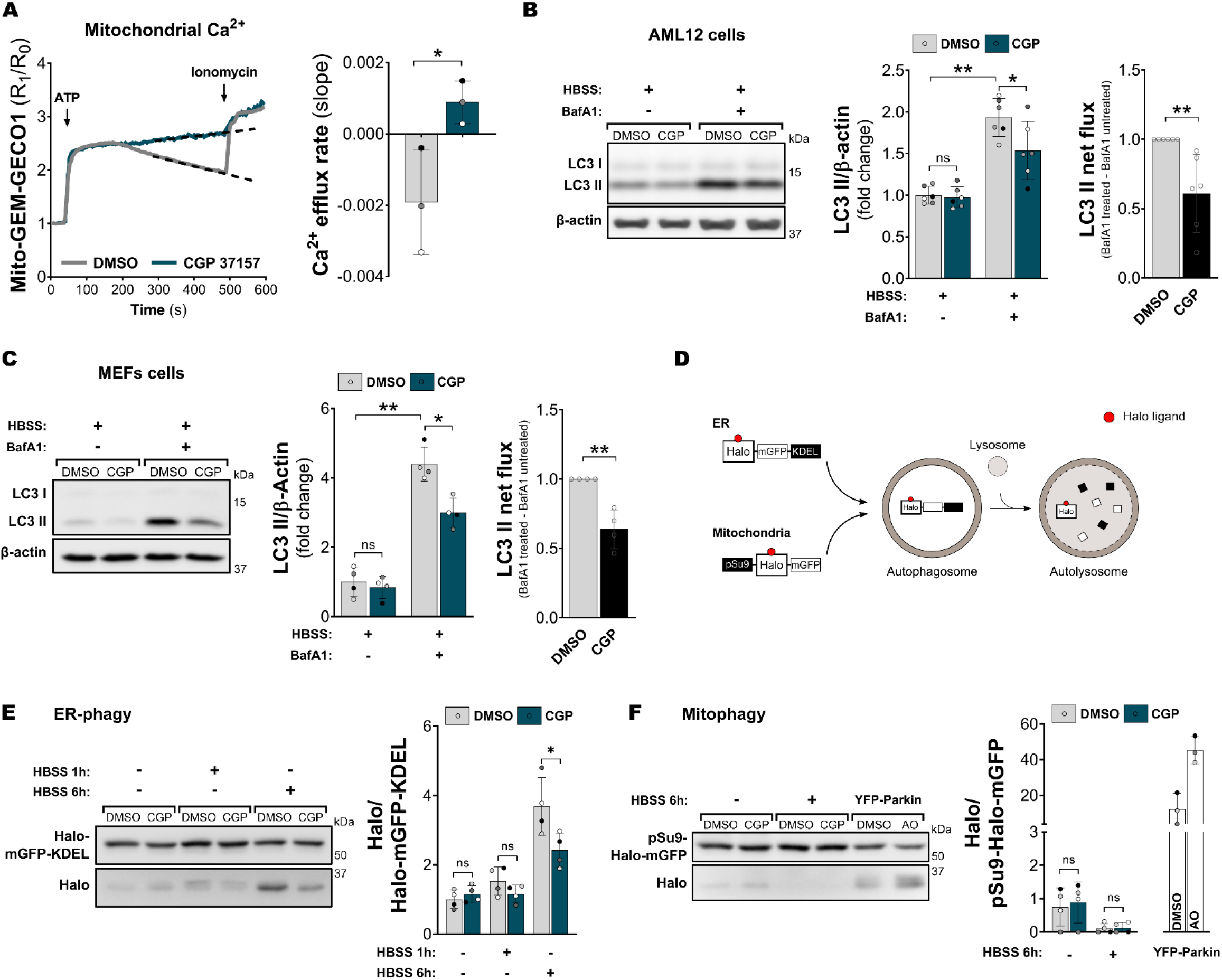
Acute NCLX inhibition decreases autophagy activation by starvation. (**A**) Measurements of intramitochondrial Ca^2+^ levels to evaluate efflux rates after 100 µM ATP stimulation in the presence of vehicle (DMSO) or 10 µM CGP 37157 (CGP) (n = 3). Data were analyzed by two-tailed paired t test. (**B**) Western blot analysis of LC3 II in AML12 cells after 1 hour incubation with HBSS in the absence or presence of Bafilomycin A1 (BafA1) during the last 30 min of treatment (n = 6). Data were analyzed by two-way RM ANOVA followed by Tukey’s multiple comparisons test. LC3 II net flux of DMSO- and CGP-treated cells were analyzed by two-tailed unpaired t test. (**C**) MEF cells; experiments performed as in B (n = 4). (**D**) Schematic representation of HaloTag-based selective autophagy assay. (**E**) ER-phagy analysis of MEF cells stably expressing the Halo-mGFP-KDEL construct after incubation in complete media (1 hour) or HBSS (1 and 6 hours) in the presence of vehicle (DMSO) or 10 µM CGP 37157 (CGP) (n = 4). Data were analyzed by two-way RM ANOVA followed by Bonferroni’s multiple comparisons test. (**F**) Mitophagy analysis of MEFs stably expressing the pSu9-Halo-mGFP construct, after incubation in complete media or HBSS for 6 h in the presence of vehicle (DMSO) or 10 µM CGP 37157 (CGP) (n = 4). Data were analyzed by two-way RM ANOVA followed by Bonferroni’s multiple comparisons test. Transient overexpression of YFP-Parking and treatment with 4 µM antimycin A plus 10 µM oligomycin (AO) was used as positive control (n = 3). Data are presented in columns with means ± SD. Dots in each group represent biological replicates, and paired biological replicates are represented by equally colored symbols. * = p < 0.05, ** = p < 0.01, *** = p < 0.001, ns = not significant.

When CGP was concomitantly added to AML12 cells during incubation with HBSS, we observed a reduction in autophagic activity, as shown by lower accumulation of LC3 II in the presence of BafA1 and lower net flux (**Figure 3B**). Interestingly, CGP also promoted the accumulation of LC3 I during autophagy induction (Fig. S3A), suggesting inhibition of the lipidation step. Moreover, inhibition of starvation-induced autophagy was observed using control versus CPG-treated mouse embryonic fibroblasts (MEFs, **Figure 3C**), showing that NCLX-dependent regulation of autophagy is not specific for hepatic cells.

Next, we addressed the question whether NCLX inhibition also affects selective autophagy. Nutrient restriction is known to favor autophagic degradation of the ER (ER-phagy) (Kohno et al., 2019, Lorenzo et al., 2022). Thus, we first measured the levels of ER-phagy in MEFs stably expressing the HaloTag-based reporter located to the ER lumen (Halo-mGFP-KDEL; Yim et al., 2022). Upon binding to its ligand, Halo becomes resistant to lysosomal degradation, while the rest of the construct is processed (**Figure 3D**; Yim et al., 2022). Therefore, the levels of Halo monomer detected by western blot are proportional to autophagic activity. Initially, we observed that treatment with CGP for 1 hour in either CM or HBSS had no impact on Halo processing (**Figure 3E**). Since incubation with HBSS for 1 hour was insufficient to induce significant ER-phagy in these cells, we extended the starvation period to 6 hours, which led to high levels of ER-phagy. Under these conditions, NCLX inhibition significantly decreased the levels of Halo monomer in comparison to DMSO (**Figure 3E**), demonstrating that starvation-induced ER-phagy is supported by NCLX activity.

Since NCLX activity is associated with mitochondrial health (Garbincius and Elrod, 2022), we also investigated whether mitophagy is affected in our model by employing a similar HaloTag-based system, now located to the mitochondrial matrix (pSu9-Halo-mGFP; Yim et al., 2022; **Figure 3F**). We validated our assay by testing the Halo processing activity in MEFs overexpressing YFP-Parkin treated with antimycin A and oligomycin (AO; **Figure 3F, white bars**, Fig S3B) as a positive control for Pink1/Parkin-dependent mitophagy activation (Lazarou et al., 2015). We found that basal mitophagy was very low in MEFs, and incubation with HBSS for 6 hours resulted in virtually undetectable levels (**Figure 3F**). This data agrees with previous findings in the literature which show protection of the mitochondrial network from degradation during upregulation of bulk autophagy by nutrient restriction (Rambold et al., 2011) and rapid degradation of selective autophagy receptor by microautophagy in this condition (Mejlvang et al., 2018). Interestingly, treatment with CGP for 6 hours had no effect on Halo processing neither in CM nor following HBSS incubation (**Figure 3F**), demonstrating that mitophagy is unaffected by NCLX activity under these conditions. Nonetheless, this does not exclude a possible role of NCLX in contexts where mitophagy levels are higher. For example, CGP was recently suggested to inhibit hypoxia-induced mitophagy (Babuharisankar et al., 2023). Additionally, another study observed increased basal mitochondrial oxidant production and slightly more mitophagosomes in colorectal cancer cells lacking NCLX (Pathak et al., 2020). This agrees with the hypothesis that NCLX activity is important to prevent mitochondrial Ca^2+^ overload in some contexts, which could trigger mitochondrial depolarization and subsequent mitophagy activation (Garbincius and Elrod, 2022). Hence, since mitophagic activity is low under our conditions, an interplay between NCLX and basal or induced mitophagy cannot be excluded and further investigation using different stimuli and cell types can provide deeper insights into this relationship.

### Autophagy inhibition is independent of changes in mitochondrial ATP and AMPK

Prior effects of NCLX in the literature have been mechanistically associated with changes in intra or extramitochondrial Ca^2+^ (Palty et al., 2010; Nita et al., 2012; Tekeuchi et al., 2013; Parnis et al., 2013; Jadiya et al., 2019; Pathak et al., 2020; Garbincius et al., 2022; Cabral-Costa et al., 2023), both of which are plausible mechanisms capable of modulating autophagic activity. In mitochondria, controlled increases in Ca^2+^ concentrations are known to promote oxidative phosphorylation, as long as they are within a specific “Goldilocks” concentration range (Vilas-Boas et al., 2023). This concentration-dependent stimulation of oxidative phosphorylation is due to the lack of activation of vital matrix metabolic enzymes when mitochondrial Ca^2+^ is too low and opening of the mitochondrial permeability transition pore (mPTP) and inhibition of respiration when Ca^2+^ levels are too high. In fact, decreased intramitochondrial Ca^2+^ caused by MCU inhibition was shown to decrease oxidative phosphorylation, and activate AMPK-dependent autophagy (Cárdenas et al., 2010). Thus, it is possible that NCLX inhibition causes the opposite effect. To address this hypothesis, we measured ATP synthesis through oxidative phosphorylation in intact AML12 cells using extracellular flux analysis. We found that neither treatment with CGP for 1 hour nor NCLX KD resulted in significant alterations in basal or ATP-linked mitochondrial oxygen consumption rates (ATP-linked OCR) (**Figure 4A**, Fig. S4A). This observation is consistent with the finding that NCLX inhibition or KD maintains intramitochondrial calcium concentrations under control conditions within the “Goldilocks” range, without impacting on oxidative phosphorylation (Vilas-Boas et al., 2023), while it still promotes changes in autophagy. As a confirmation that NCLX effects over autophagy are not due to changes in ATP production, we observed that CGP was able to proportionally inhibit starvation-induced autophagy in MEFs lacking AMPK expression (AMPKα1/2 KO) (**Figure 4B**), proving that this effect is AMPK-independent.

**Figure 4.**
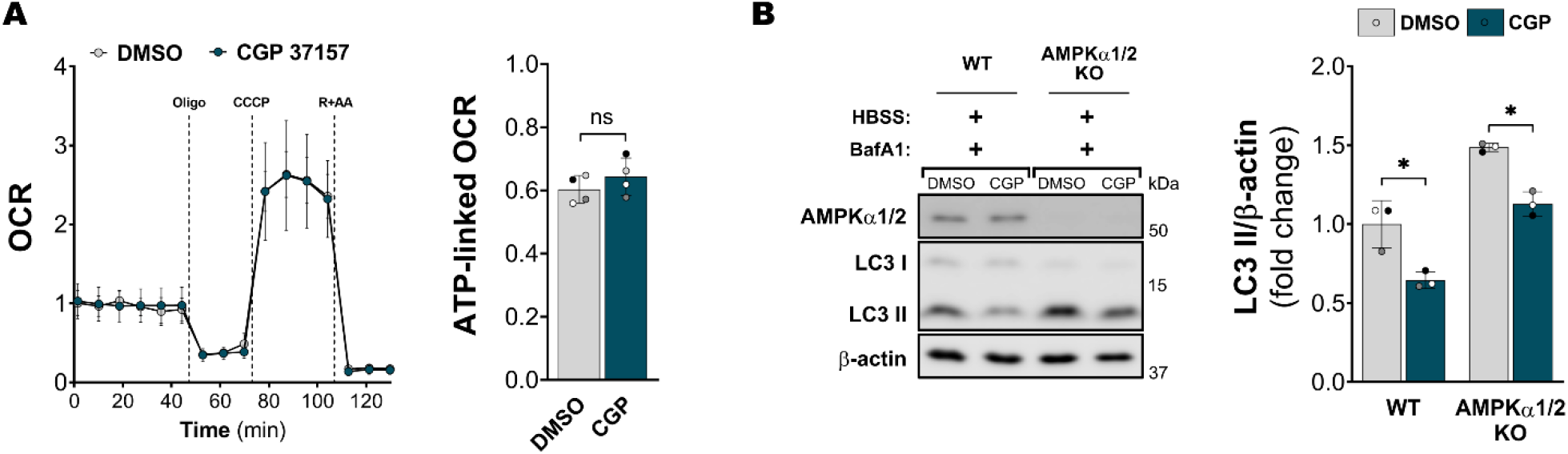
Autophagy inhibition by CGP 37157 is independent of changes in mitochondrial ATP and AMPK. (**A**) Oxygen consumption rates (OCR) linked to ATP synthesis (oligomycin-sensitive respiration) of AML12 cells treated with either DMSO or CGP (n = 4). Data were analyzed by two- tailed paired t test. (**B**) Western blot analysis of LC3 II in wild-type (WT) or knockout for AMPKα1/2 (AMPKα1/2 KO) MEFs after 1 hour incubation with HBSS plus Bafilomycin A1 (BafA1), in the presence of vehicle (DMSO) or 10 µM CGP 37157 (CGP) (n = 3). Data were analyzed by two-way RM ANOVA followed by Tukey’s multiple comparisons test. Data are presented in columns with means ± SD. Dots in each group represent biological replicates, and paired biological replicates are represented by equally colored symbols. * = p < 0.05, ** = p < 0.01, *** = p < 0.001, ns = not significant.

### NCLX inhibition acts on early steps of autophagy

Since our previous data suggested that NCLX activity regulates autophagosome formation in MEFs and AML12 cells, we decided to investigate the formation of the early autophagy machinery puncta, typically observed in response to amino acid starvation. We immunostained MEFs for FIP200, which marks the formation of the ULK1 complex in autophagy initiation sites, and for ATG16L1, which marks the phagophore (Zachari and Ganley, 2017). As expected, cells incubated with HBSS for 1 hour showed increased FIP200 and ATG16L1 puncta formation (**Figure 5**). Conversely, the presence of CGP significantly inhibited puncta formation detected using both markers (**Figure 5**), showing that NCLX activity supports the recruitment of autophagy initiation machinery during starvation.

**Figure 5.**
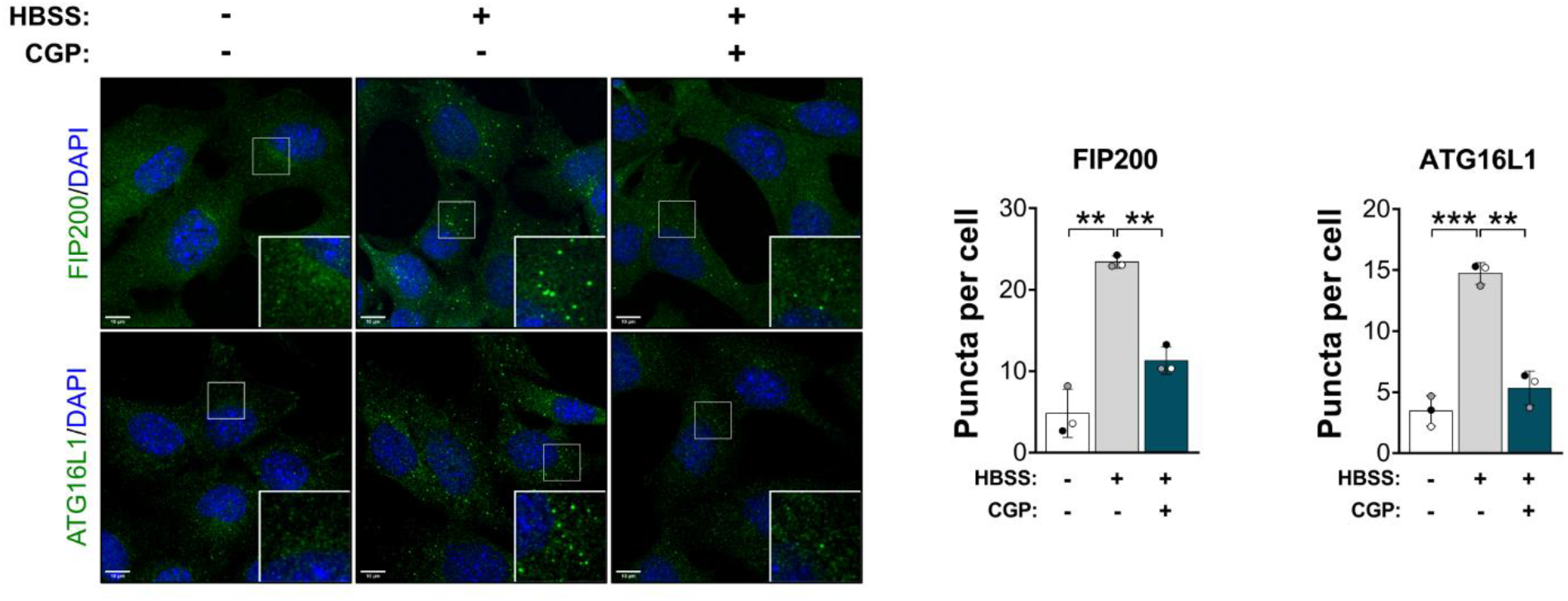
NCLX inhibition affects early steps of autophagy activation. Immunofluorescence analysis of FIP200 and ATG16L1 puncta formation in MEF cells incubated with complete media or 1 hour of HBSS in the presence of vehicle (DMSO) or 10 µM CGP 37157 (CGP) (n = 3). Data were analyzed by one-way RM ANOVA followed by Bonferroni’s multiple comparisons test. Data are presented in columns with means ± SD. Dots in each group represent biological replicates, and paired biological replicates are represented by equally colored symbols. * = p < 0.05, ** = p < 0.01, *** = p < 0.001, ns = not significant.

### Impaired autophagy promoted by NCLX inactivation is the result of reduced intracellular Ca^2+^

In addition to the accumulation of Ca^2+^ in the mitochondrial matrix, decreased Ca^2+^ release into the cytosol is a direct downstream effect of NCLX inhibition (Palty et al., 2010; Cabral-Costa et al., 2023). Together with the ER and plasma membrane, mitochondrial Ca^2+^ uptake is responsible for Ca^2+^ clearance from the cytosol during signaling events (Rizzuto et al., 2012). Interestingly, early steps of autophagy activation during starvation were shown to be regulated by Ca^2+^ signaling. For instance, a recent study has shown that starvation-induced subcellular increases in cytosolic Ca^2+^ trigger FIP200 puncta formation in autophagosome formation sites. Considering our observation that treatment with CGP inhibited FIP200 puncta formation, we questioned if NCLX was regulating cytosolic Ca^2+^ levels in our model. These were assessed through Fura-2 fluorescence ratios upon stimulation with 100 µM ATP and in the presence of DMSO (control) or CGP (**Figure 6A**). Interestingly, although acute inhibition of NCLX did not affect the ATP-induced Ca^2+^ peak (**Figure 6A**), clearance of Ca^2+^ from the cytosol was faster, as shown by a substantial reduction in the half-life decay and the area under the curve (AUC) (**Figure 6A**). Indeed, decreased cytoplasmic Ca^2+^ levels due to NCLX inhibition have been observed in prior studies (Parnis et al., 2013; Cabral-Costa et al., 2023). Consistently, siNCLX cells presented decreased Ca^2+^ release after ATP stimulus (Fig S5A, Nita et al., 2012; Stavsky et al., 2012; Kim et al., 2012; Parnis et al., 2013). In KD cells, the half-life decay was strikingly higher (Fig S5A), indicating that other Ca^2+^ clearance pathways are strongly affected by decreased NCLX expression, and further linking reduced NCLX activity to impaired cytoplasmic Ca^2+^ signals. Indeed, NCLX KD has been extensively associated with decreased cellular Ca^2+^ entry and, consequently, low Ca^2+^ levels in the ER (Takeuchi et al., 2013; Parnis et al., 2013; Ben-Kasus Nissim et al., 2017; Emrich et al., 2022; Takeuchi and Matsuoka, 2022).

**Figure 6.**
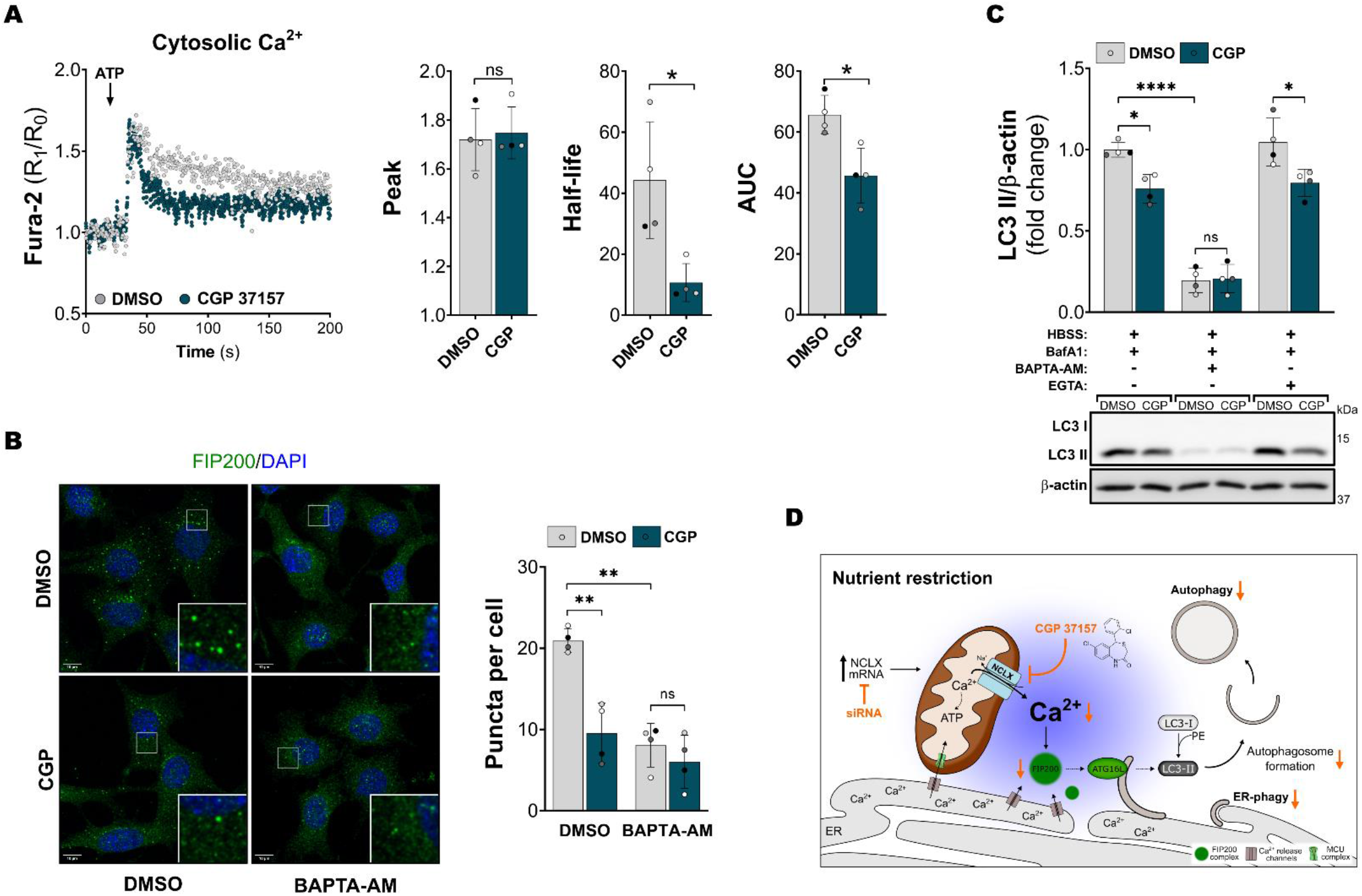
Impaired autophagy is a result of reduced cytoplasmic Ca^2+^. (**A**) Cytosolic Ca^2+^ measurements using Fura-2 AM after stimulation with 100 µM ATP; bar graphs showing quantifications of Ca^2+^ peaks, the half-life of one phase decay after the peak and area under the curve (AUC) (n = 4). Data were analyzed by two-tailed paired t test. (**B**) FIP200 puncta formation in MEFs pre-incubated with either DMSO or 50 µM BAPTA-AM for 1 hour prior to incubation with HBSS for 1 hour in the presence either DMSO or 10 µM CGP 37157 (CGP) (n = 4). (**C**) Western blots for LC3 II in MEFs incubated with HBSS plus BafilomycinA1 (BafA1) for 1 hour in the presence either DMSO or 10 µM CGP 37157 (CGP). 50 µM BAPTA-AM was pre-incubated for 1 hour and 5 mM EGTA was incubated with HBSS (n = 4). Data were analyzed by two-way RM ANOVA followed by Tukey’s multiple comparisons test. Data are presented in columns with means or means ± SD. Dots in each group represent biological replicates, and paired biological replicates are represented by equally colored symbols. * = p < 0.05, ** = p < 0.01, *** = p < 0.001, **** = p < 0.0001, ns = not significant. (**D**) Schematic representation of our hypothesis over NCLX regulation of autophagy.

To test whether the changes in intracellular Ca^2+^ homeostasis could lead to defective autophagy, we pre-incubated cells for 1 hour with the cell-permeant Ca^2+^ chelator BAPTA-AM before incubation with HBSS. As expected, pre-incubation with BAPTA-AM prevented FIP200 puncta formation by HBSS (**Figure 6B**). Consequently, it also resulted in decreased LC3 II accumulation in the presence of BafA (**Figure 6C**, Fig 56B) and accumulation of LC3 I (Fig S5B), similar to the observations with NCLX inhibition. To test if NCLX regulation of autophagy requires intracellular Ca^2+^ signaling, we verified the concomitant effects of CGP and BAPTA-AM on FIP200 puncta formation and LC3 II levels and did not observe any additive effect of CGP in cells pre-incubated with the Ca^2+^ chelator (**Figure 6B**, **6C,** Fig S5B). On the other hand, CGP caused autophagy inhibition when extracellular Ca^2+^ was chelated by the presence of cell-impermeable EGTA (**Figure 6C**), thus excluding possible unspecific effects of CGP on Ca^2+^ transport of the plasma membrane. Although we propose that NCLX effects on autophagy initiation are due to the reduction of cytosolic Ca^2+^ levels, BAPTA-AM may also deplete Ca^2+^ within the mitochondria, so we cannot exclude the possibility of other unknown mechanisms where intramitochondrial Ca^2+^ could be affecting autophagy activation without changes in ATP production. Irrespectively, these results clearly demonstrate that mitochondrial Ca^2+^ efflux through NCLX is involved in Ca^2+^ signaling events required for the early steps of autophagy activation.

Intracellular Ca^2+^ signals generated during amino acid and serum starvation are known to be required for autophagy activation (Ghislat et al., 2012). However, the origin of this Ca^2+^ and its downstream effects are multiple and complex. Elevation in cytoplasmic Ca^2+^ has been proposed to activate autophagy through the CaMKKI-β-AMPK-mTORC1 pathway (Høyer-Hansen et al., 2007; Ghislat et al., 2012). However, our results demonstrating that the CGP effect is AMPK-independent support the idea that other steps of autophagy machinery are being affected. In line with this, transient increases in Ca^2+^ concentrations generated on the ER or lysosome surface trigger liquid-like FIP200 puncta formation. (Zheng et al., 2022). In agreement, we demonstrate here that Ca^2+^ efflux from mitochondria through NCLX also regulates FIP200 puncta formation during starvation. Consistently, ER-mitochondria contact sites have been implicated as sources of an autophagosomal membrane (Hamasaki et al., 2013).

The schematic summary of our hypothesis regarding the interplay between NCLX and autophagy is shown in **Figure 6D**. Our findings add to the current knowledge by showing that NCLX is both regulated by autophagy-inducing nutrient deprivation and contributes to autophagy activation, as seen by both silencing and acute pharmacological inhibition experiments. These results indicate that NCLX is an important new regulatory node, integrating autophagy control by Ca^2+^ and cellular responses to nutrient availability.

## Methods

### Chemicals, reagents and antibodies

siRNA molecules, insulin-transferrin-selenium (ITS) solution, fetal bovine serum (FBS), DMEM/F-12, high glucose DMEM, HBSS, Ripa Buffer, Lipofectamine^™^ RNAiMAX, Lipofectamine^™^ 2000, Opti-MEM^™^, High-Capacity cDNA Reverse Transcription Kit, Platinum^™^ SYBR^™^ Green qPCR SuperMix-UDG, protease and phosphatase inhibitor cocktails, Pierce*™* BCA Protein Assay Kit, Fura-2 AM, Ca^2+^ Green^™^-5N, and ProLong^™^ Diamond were purchased from Thermo Fisher Scientific (Waltham, MA, USA). CGP 37157 (1114) was purchased from Tocris/Bio-Techne (Bristol, UK). BAPTA-AM (A1076) and Bafilomycin A1 (SML1661) were purchased from Sigma-Aldrich (St. Louis, MO, USA). Ionomycin (11932) was purchased from Cayman Chemical (Ann Arbor, MI, USA).

Primary antibodies against β-actin (8226; 1:5,000) were purchased from Abcam (Cambridge, UK). Primary antibodies against LC3 A/B (12741; 1:1,000), LC3B (2775; 1:1,000), Parkin (4211; 1:1000) and α-tubulin (3873; 1:10,000) were purchased from Cell Signaling Technology (Danvers, MA, USA). NCLX antibody (orb317904; 1:500) was purchased from Biorbyt (Cambridge, UK). Primary antibody against HaloTag^®^ (G9211; 1:1000) was purchased from Promega (UK).

### Caloric restriction diet and mitochondrial isolation

Eight-week-old male Swiss mice were divided randomly into *ad libitum* (AL) and caloric restriction (CR) groups, eating 40% less calories than AL animals, with micronutrient supplementation, for 4 weeks (Menezes-Filho et al., 2017). The diet was adjusted weekly according to the AL animals’ food intake. Both chow diets were prepared by Rhoster^®^ (Campinas, SP, Brazil). All animal experiments were approved by the animal ethical committee and followed National Institutes of Health guidelines for animal welfare.

After the diet period, mitochondria were isolated from the livers of mice as described previously (Serna et al., 2022). Briefly, the livers were minced, washed repeatedly with ice-cold PBS and suspended in isolation buffer (250 mM sucrose, 10 mM HEPES, 1 mM EGTA, 1 mM EDTA, 1 mg/ml BSA fatty acid-free). Tissues were homogenized using a Potter-Elvehjem tissue grinder, and mitochondria were isolated by differential centrifugation.

### Cell cultures

AML12 cells were purchased from ATCC^®^ and cultivated in complete media (CM) consisting of DMEM/F-12 media supplemented with 10% FBS (v/v), 1% antibiotics (100 U/ml penicillin, 0.1 mg/ml streptomycin), and 1% ITS (Gibco^™^, 41400045). HEK 293FT cells (kindly donated by Prof. Alexandre-Bruni Cardoso, Universidade de São Paulo), MEFs (a gift from Prof. Noboru Mizushima, University of Tokyo; Kataura et al., 2022), WT and AMPKα1/2 KO MEFs (a gift from Prof. Benoit Viollet, Institut National de la Santé et de la Recherche Médicale) were cultivated in High Glucose DMEM media supplemented with 10% FBS (v/v) and 1% antibiotics. Cells were kept in a humidified incubator with a 5% CO_2_ atmosphere, at 37°C, and trypsinized every 48-72 hours. The passage number was controlled up to 35. HBSS (Gibco^™^, 14175095) supplemented with 120 mg/L CaCl_2_, 30 mg/L MgCl_2_, and 50 mg/L MgSO_4_ was used for serum and amino acid starvation experiments.

### Small interference RNA (siRNA) transfection

AML12 cells were transfected with siRNAs targeting NCLX (Ambion^®^, ID s100747) or with a Negative Control (NC) sequence (Ambion^®^, 4390843) at a final concentration of 20 nM. Transfections were performed using the Lipofectamine^™^ RNAiMAX (Invitrogen^™^, 13778030) reagent through a reverse transfection protocol. Briefly, Lipofectamine^™^ and siRNAs were mixed in Opti-MEM^™^ media and incubated for 5-10 minutes at room temperature. Cells were suspended in transfection media (DMEM/F-12, 10% FBS, 1% ITS, without antibiotics) and added to the plate well on top of the Lipofectamine^™^-siRNA complexes. The transfection media was replaced by CM 24 hours later, and experiments were performed 48-72 hours after transfection. For experiments performed at 72 hours, the cells were trypsinized and re-seeded in new culture plates 24 hours earlier.

### RT-qPCR

RNA was isolated from cells using Trizol^®^ reagent following the manufacturer’s instructions. The concentration of isolated RNA was quantified using a NanoDrop^®^ spectrophotometer and 1-2 µg was used for cDNA synthesis with a High-Capacity cDNA Reverse Transcription Kit (Applied Biosystems^™^, 4368814). qPCR reactions were conducted using a 7500 Real-Time PCR System (Applied Biosystems^™^) with Platinum^™^ SYBR^™^ Green qPCR SuperMix-UDG (Invitrogen^™^, 11733046), 10 ng of cDNA, and the specific primers described in Supplementary Table 1. The Ct values were analyzed through the 2^(-ΔΔCt)^ method and the housekeeping gene or combination of genes was chosen by testing the expression of *Hmbs*, *B2m* and *Actb* in each experimental group. The compared stability value was calculated using NormFinder Software (Andersen et al., 2004). The mean Ct value between *Hmbs* and *B2m* expression was used as housekeeping for experiments containing siRNA-mediated knockdown. For experiments containing HBSS time curve treatments, a *B2m* Ct value was used.

### Western blots

Cells were washed with 1X ice-cold PBS and lysed in RIPA buffer containing protease and phosphate inhibitor cocktail (Thermo Scientific^™^, 78443). Cell lysates were sonicated using a probe sonicator and protein concentration was quantified using the Pierce*™* BCA Protein Assay Kit (Thermo Scientific^™^, 23227). Samples were prepared in 1X Laemmli buffer (2% SDS, 10% glycerol, 0.062 M Tris pH 6.8, 0.002% bromophenol blue, 5% 2-mercaptoethanol), loaded onto SDS-PAGE for gel electrophoresis separation, and transferred overnight to PVDF membranes using a wet transfer system. Membranes were blocked in TTBS (20 mM Tris pH 7.5, 150 mM NaCl and 0.1% Tween 20) containing 3-5% BSA (w/v) before overnight incubation with primary antibodies at 4°C. After washing with TTBS, membranes were incubated with fluorescent IRDye^®^ secondary antibodies (1:15,000) for 1 hour at room temperature. Fluorescence was detected using the ChemiDoc^™^ Imaging System (Bio-Rad) and band densitometry was quantified using Fiji ImageJ software.

### mCherry-EGFP-LC3B expression, imaging, and analysis

To evaluate autophagic activity, we generated an AML12 cell line stably expressing mCherry- EGFP-tagged LC3B protein. Retroviruses were produced by transfecting HEK 293FT cells with a pBABE-puro-mCherry-EGFP-LC3B plasmid (Addgene, 22418; gifted by Jayanta Debnath, N’Diaye et al., 2009) and a PCL-Eco packaging plasmid (Addgene, 12371; gifted by Inder Verma, Naviaux et al., 1996). 48-72 hours after transfection, the media from the HEK 293FT cells was used to transduce AML12 cells through multiple rounds of infection. 24 hours after the last infection, cells were kept in CM containing 10 µg/mL puromycin for five days. After selection, cells were cultivated in CM supplemented with 1 µg/mL puromycin, which was removed from the media during experiments.

For experiments, AML12-LC3 cells were transfected with specific siRNAs using the same protocol described in the “*Small interference RNA (siRNA) transfection”* section. 48 hours after transfection, cells were trypsinized, seeded on poly-D-lysine-coated 12 mm coverslips (Thorlabs, CG15NH) and incubated for an additional 24 hours before treatments. After treatments, media was removed, and cells were fixed in 4% paraformaldehyde for 15 minutes at room temperature. Coverslips were washed once with 1X PBS containing 0.1 M Glycine, washed two times with 1X PBS and incubated with DAPI (0.1 µg/mL) for 10 minutes at room temperature. Finally, coverslips were washed three times with 1X PBS and mounted onto glass slides using ProLong^™^ Diamond Antifade Mountant (Invitrogen^™^, P36965). Images were acquired using a Leica DMi8 microscope with an HC PL APO CS2 63x/1.40 OIL UV objective. At least 100 cells per group were imaged and each channel was acquired through 2 µm on the Z-axis dimension (total of 6 steps). Images were processed using the Leica 3D BlindDeblur deconvolution tool and mCherry^+^/GFP^+^ (autophagosome), mCherry^+^/GFP^-^ (autolysosome), and cell numbers (DAPI) were quantified using Fiji ImageJ software.

### Proteasome activity

72 hours after siRNA transfection, cells were trypsinized, washed once with 1X cold PBS and resuspended in experimental buffer (5 mM MgCl_2_, 50 mM Tris pH 7.5). Next, cells were lysed using an insulin syringe and centrifuged for 10 min at 10000 × g, at 4°C for debris separation. Protein concentration was quantified using the Pierce*™* BCA Protein Assay Kit (Thermo Scientific^™^, 23227) and 30 µg of protein (in 100 µL of experimental buffer) was added to each well for the assay. To measure proteasome (chymotrypsin-like) activity, cell extracts were incubated with 250 µM fluorogenic peptide Suc-LLVY-AMC and the increase in AMC fluorescence was monitored for 10 minutes at 37°C, using a Flex Station 3 (Molecular Devices) microplate fluorescence spectrophotometer (λex = 341 nm, λem = 441 nm). The slope of fluorescence increase over time was quantified using GraphPad Prism 7 software.

### Mitochondrial Ca^2+^ efflux in intact cells

To estimate mitochondrial Ca^2+^ efflux rates in live cells, we measured intramitochondrial free Ca^2+^ levels by transfecting AML12 cells with the CMV-mito-GEM-GECO1 plasmid (Addgene, 32461; gifted by Robert Campbell, Zhao et al., 2011). Briefly, cells were seeded on a glass bottom culture dish (Greiner Bio-One, 627870) and, 24 hours later, the media was replaced with transfection media (DMEM/F-12, 10% FBS, 1% ITS, without antibiotics). The plasmid was transfected using Lipofectamine^™^ 2000 Transfection Reagent (Invitrogen^™^, 11668019) at a 1:1 µg DNA to µL Lipofectamine^™^ ratio, following the manufacturer’s protocol. The media was replaced for CM 24 hours after transfection and the cells were imaged after an additional 24 hours. For imaging, the media was replaced for HBSS (5.5 mM D-glucose, 120 mg/L CaCl_2_, 30 mg/L MgCl_2_, 50 mg/L MgSO_4_, 60 mg/L KH_2_PO_4_, 48 mg/L Na_2_HPO_4_, 400 mg/L KCl, 8 g/L NaCl, 350 mg/L NaHCO_3_, 5 mM HEPES) containing either DMSO or CGP 37157 and fluorescence was acquired in a Zeiss LSM 780-NLO confocal microscope using a 405 nm diode laser and a C-Apochromat” 63x/1.20 DIC objective, in an environment kept at 37°C, with 5% CO_2_ atmosphere. During acquisition, cells were imaged every 4 seconds for at least 15 minutes. 100 µM ATP was added to stimulate intramitochondrial Ca^2+^ increases and 10 µM ionomycin was used to compare the maximum fluorescence ratio between groups. Analysis was performed using the Fiji ImageJ software and results are shown as the ratio between emissions at 455 nm and 511 nm relative to the basal ratio (R_1_/R_0_). Mitochondrial efflux rates were calculated as the slope after the ATP-stimulated Ca^2+^ peak, using the GraphPad Prism 7 software.

### HaloTag-based selective autophagy assay

To measure ER-phagy and mitophagy levels, we generated MEFs stably expressing the HaloTag7-mGFP-KDEL or pSu9-HaloTag7-mGFP constructs, respectively. For that, retroviruses were produced by using Lipofectamine^™^ 2000 Transfection Reagent (Invitrogen^™^, 11668019) to co-transfect HEK 293FT cells with plasmids containing the packaging gag/pol and envelope pCMV- 435VSV-G genes and either the pMRX-IB-HaloTag7-mGFP-KDEL (Addgene, 184904; gifted by Noboru Mizushima, Yim et al., 2022) or the pMRX-IB-pSu9-HaloTag7-mGFP (Addgene, 184905; gifted by Noboru Mizushima, Yim et al., 2022). After 293FT cell transfection (48 h), the media was collected and viral particles were concentrated using Lenti-X™ Concentrator (Takara Bio Inc., 631231). For transduction, MEFs were seeded in a 6 well culture plate and, 24 hours later, 1 µL of concentrated viral particles was added in 1 mL of CM containing 10 µg/mL of polybrene. On the next day, the media was replaced with selection media containing 8 µg/mL of blasticidin, where the cells were kept for 48 hours. After selection, cells were cultivated in CM supplemented with 2 µg/mL blasticidin, which was removed during experiments.

For ER-phagy experiments, 1.2×10^5^ cells/well were seeded on a 6 well plate. 48 hours later, 30 µg/mL of 7-bromo-1-heptanol (HaloTag-blocking agent/ligand) was added to the cells for 30 minutes prior to treatments. For treatments, cells were washed twice with PBS and incubated under conditions described. For mitophagy experiments, 7.5×10^4^ cells were seeded on 6-well plates and treated 72 hours later. For positive control experiments, 1.5×10^5^ cells/well were seeded concomitantly on 6-well plates and, 24 hours later, each well was transfected with 1.5 µg of the plasmid containing the YFP-Parking gene (Addgene, 23955; gifted by Richard Youle, Narendra et al., 2008) and 3 µL of Lipofectamine^™^ 2000 Transfection Reagent (Invitrogen^™^, 11668019), according to manufacturer instructions. 48 hours after transfection and 72 hours after seeding, all cells were treated as described in media containing 30 µg/mL of 7-Bromo-1-heptanol, which was present during the full incubation time. Treatment with 4 µg/mL antimycin A and 10 µg/mL oligomycin (AO) was used to stimulate mitophagy in YFP-Parkin cells. In both experiments, the cells were collected in RIPA buffer after treatment periods and processed as described in “*Western blot”* section using Halo primary antibody. ER-phagy or mitophagy levels were analyzed as quantification of Halo monomer/Full-length construct.

### Confocal immunofluorescence

For the visualization of FIP200 and ATG16L1 puncta, MEF cells were immunostained as follows. Briefly, cells were seeded onto 13 mm glass coverslips and treated as indicated. After treatments, the media was removed, and cells were fixed with 4% formaldehyde for 10 minutes at room temperature. Next, coverslips were washed with PBS and cells were permeabilized with a PBS solution containing 0.5% Triton X-100 for 10 min. Cells were then washed 3 times with PBS and incubated with blocking solution (PBS, 5% Normal Goat Serum, 0.05% Tween) for 1 hour at room temperature. Primary antibodies for FIP200 (Cell Signaling Technology, 12436S;1:100) and ATG16L1 (Cell Signaling Technology, 8089S; 1:100) were incubated overnight at 4°C. Cells were washed 3 times with PBS containing 0,05% Tween (PBS-T), incubated for 1 hour at room temperature with secondary antibodies (Alexa Fluor^®^ 488, 1:1000) and washed again 3 times with PBS-T. Coverslips were mounted using ProLong^™^ Gold Antifade Mountant with DAPI (Invitrogen^™^, P36935). Images were acquired using a Zeiss LSM700 confocal microscope with a Plan-Apochromat 63x/1.40 Oil DIC M27 objective. Quantification of puncta number per cell was performed using Fiji ImageJ software.

### ATP-linked respiration

Mitochondrial oxygen consumption linked to ATP production was measured using a Seahorse Extracellular Flux Analyzer (Agilent Technologies, Santa Clara, CA, EUA) as described previously (Vilas-Boas et al., 2023). Briefly, 3×10^4^ cells per well were seeded in CM 24 hours before analysis. On the next day, cells were washed three times with DMEM/F-12 media supplemented with 1% antibiotics and 5 mM HEPES (without bicarbonate and FBS) and were incubated in the same media for 1 hour in the absence or presence of 10 µM CGP 37157, at 37°C in a normal atmosphere. Oxygen consumption rates were measured under basal conditions and after the injection of 1 µM oligomycin to calculate ATP-linked respiration. After the experiment, cells were lysed in Ripa Buffer and total protein was quantified using the Pierce*™* BCA Protein Assay Kit (Thermo Scientific^™^, 23227) for data normalization.

### Cytosolic Ca^2+^ measurements

Initially, cells were incubated with a mix of 5 µM Fura-2 AM (Invitrogen^™^, F1221) and 0.025% Pluronic® F-127 (diluted in CM) for 10 minutes at 37°C for Fura-2 loading. Next, cells were trypsinized, centrifuged, washed once with 1X PBS, and resuspended in HBSS (5.5 mM D-Glucose, 120 mg/L CaCl_2_, 30 mg/L MgCl_2_, 50 mg/L MgSO_4_, 60 mg/L KH_2_PO_4_, 48 mg/L Na_2_HPO_4_, 400 mg/L KCl, 8 g/L NaCl, 350 mg/L NaHCO_3_, 5 mM HEPES). The cell number was determined and 1×10^6^cells were added to a cuvette to a final volume of 2 mL. Importantly, for experiments using pharmacological inhibition, DMSO (equal volume, as a control) or CGP 37157 were added to the cuvette before the fluorescence measurement started. The cuvette was placed in a F4500 Hitachi Fluorimeter at 37°C with continuous stirring and Fura-2 fluorescence was detected at λex = 340/380 nm and λem = 505 nm. The addition of 100 µM ATP was used to stimulate an increase in cytosolic Ca^2+^ levels, which were analyzed as the ratio between fluorescence emission at 340 nm and 380 nm excitation (340/380) relative to the basal ratio (R_1_/R_0_). Peak height, area under the curve (AUC) and half-life after the peak were calculated using GraphPad Prism 7 software.

### Statistical analysis

Statistical analysis was performed using GraphPad Prism 7 software. At least three independent biological replicates were conducted for each experiment, and these are represented by individual dots in the graphs. Dots of the same color/pattern in different groups represent paired biological replicates. Data comparing two groups were analyzed with Student’s t-test. Data comparing three different groups were analyzed with one-way randomized blocks ANOVA followed by Bonferroni’s multiple comparison test. Data comparing two factors (e.g., siRNA and Bafilomycin A1) were analyzed with two-way randomized blocks ANOVA followed by either Bonferroni’s (when comparing only column effects) or Tukey’s (when comparing rows and column effects) multiple comparisons test. Results are shown as means ± SD and * = p < 0.05, ** = p < 0.01, *** = p < 0.001, **** = p < 0.0001, ns = not significant.

## Acknowledgements

This work was supported by the Fundação de Amparo à Pesquisa do Estado de São Paulo (FAPESP) under grant numbers 13/07937-8, 20/06970-5, Conselho Nacional de Desenvolvimento Científico e Tecnológico (CNPq), and Coordenação de Aperfeiçoamento de Pessoal de Nível Superior (CAPES) line 001 to A.J.K. V.M.R., J.D.S., E.A.V.B., F.M.C and A.J.K. were supported by FAPESP fellowships #2019/18402-4, #2022/08581-1, #2019/05226-3, #2021/02481-2, #2017/14713-0. T.K. was supported by Fellowships from Uehara Memorial Foundation and the International Medical Research Foundation. V.I.K. acknowledges grants from BBSRC (BB/M023389/1 and BB/R008167/2), and RESETageing H2020 grant (952266). We acknowledge Silvânia M. P. Neves and her animal facility crew for exceptional expert animal care; Natália Soares Ferreira and Mario Costa Cruz from the *Centro de Facilidades de Apoio à Pesquisa* (CEFAP) for excellent technical assistance in the Confocal experiments; Camille C. Caldeira da Silva and Sirlei Mendes de Oliveira for the remarkable technical support. We also acknowledge Dr. Sergio Menezes-Filho for initial animal experiments and Prof Alexandre Bruni-Cardoso for reagent donation. Parts of Figure 6D were drawn using pictures from Servier Medical Art. Servier Medical Art by Servier is licensed under a Creative Commons Attribution 3.0 Unported License (https://creativecommons.org/licenses/by/3.0/).

## Author contributions

V.M.R., V.I.K. T.K., and A.J.K. designed the research project; all authors designed and discussed experiments; V.M.R., J.D.C.S., E.A.V.B., J.V.C.C and T.K. analyzed data; V.M.R., J.D.C.S., E.A.V.B., J.V.C.C., F.M.C. and T.K. performed research; V.M.R. and A.J.K. primarily wrote the paper, which was perfected, revised and approved by all authors.

## Data availability

Full datasets are presented in the manuscript, with individual biological repetitions included as the distinct types of individual datapoint symbols. Datasheets of time traces, spreadsheets and images are available upon request to the authors and will be deposited in the institutional University of São Paulo data repository (persistent link requested and pending).

---

**Extended Figure 1.**
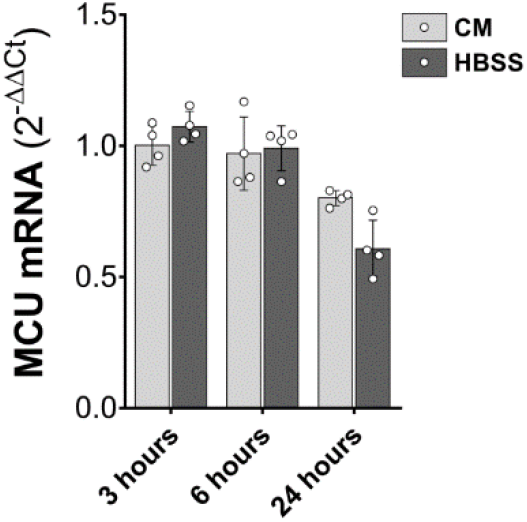
RT-qPCR for MCU mRNA in AML12 cells incubated with either complete media (CM) or HBSS for 3, 6, and 24 hours (n = 4).

**Extended Figure 2.**
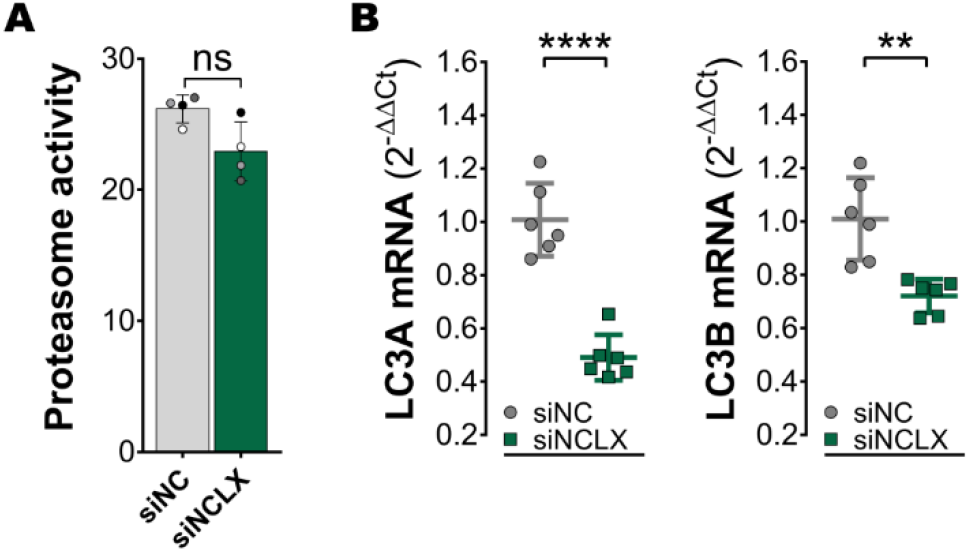
AML12 cells transfected with NCLX (siNCLX) or negative control (siNC) siRNAs levels of (**A**) proteasome activity (n = 4) and (**B**) LC3A and LC3B mRNA analyzed by RT-qPCR (n = 6). Data were analyzed by two-tailed paired t test. Data are presented in columns with means ± SD. * = p < 0.05, ** = p < 0.01, *** = p < 0.001, ns = not significant.

**Extended Figure 3.**
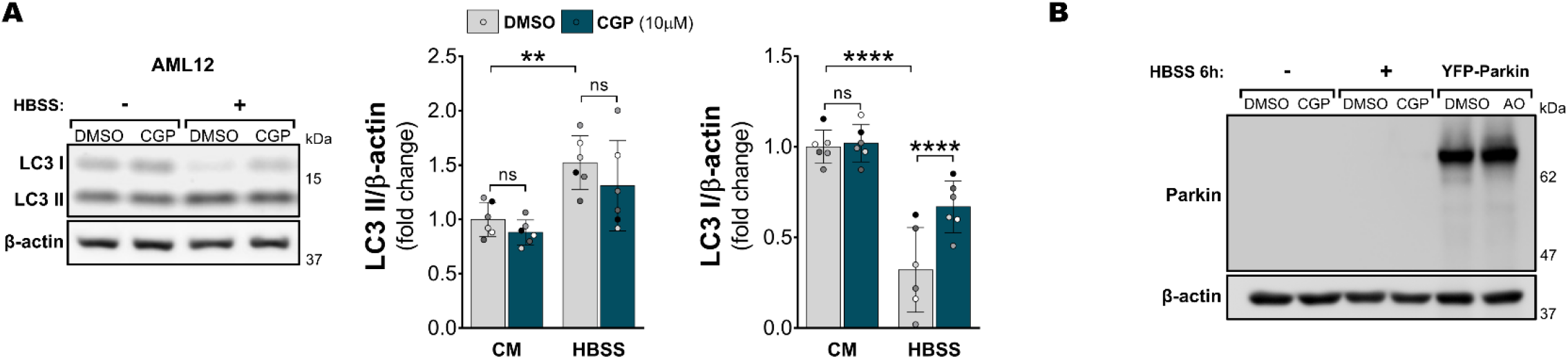
(**A**) Western blots for LC3 II and LC3 I of AML12 cells after 1 hour incubation with either complete media (CM) or HBSS in the presence of either DMSO or CGP (n = 6). Data were analyzed by two-way RM ANOVA followed by Tukey’s multiple comparisons test. Data are presented in columns with means ± SD. * = p < 0.05, ** = p < 0.01, *** = p < 0.001, ns = not significant. (**B**) Representative Parkin western blot analysis of MEFs stably expressing pSu9-Halo-mGFP showing non-detectable levels of endogenous Parkin and YFP-Parkin overexpression.

**Extended Figure 4.**
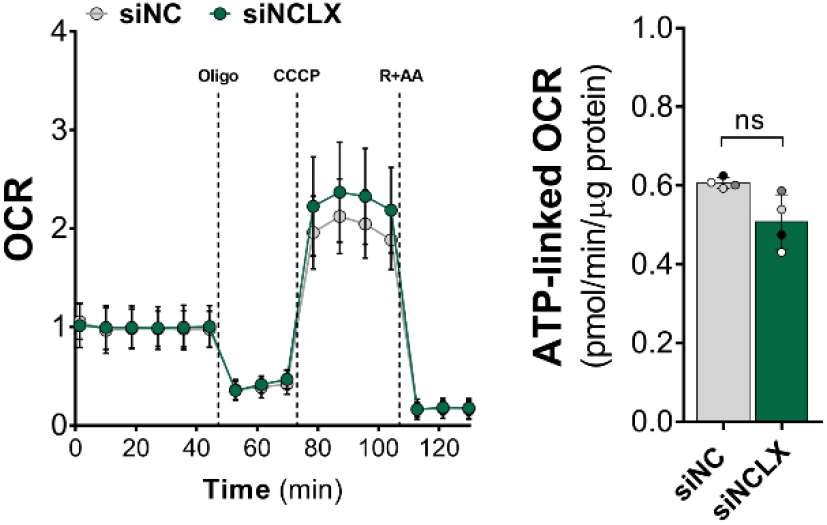
Oxygen consumption rates (OCR) linked to ATP synthesis (oligomycin-sensitive respiration) of AML12 cells transfected with NCLX (siNCLX) or negative control (siNC) siRNAs (n = 4).

**Extended Figure 5.**
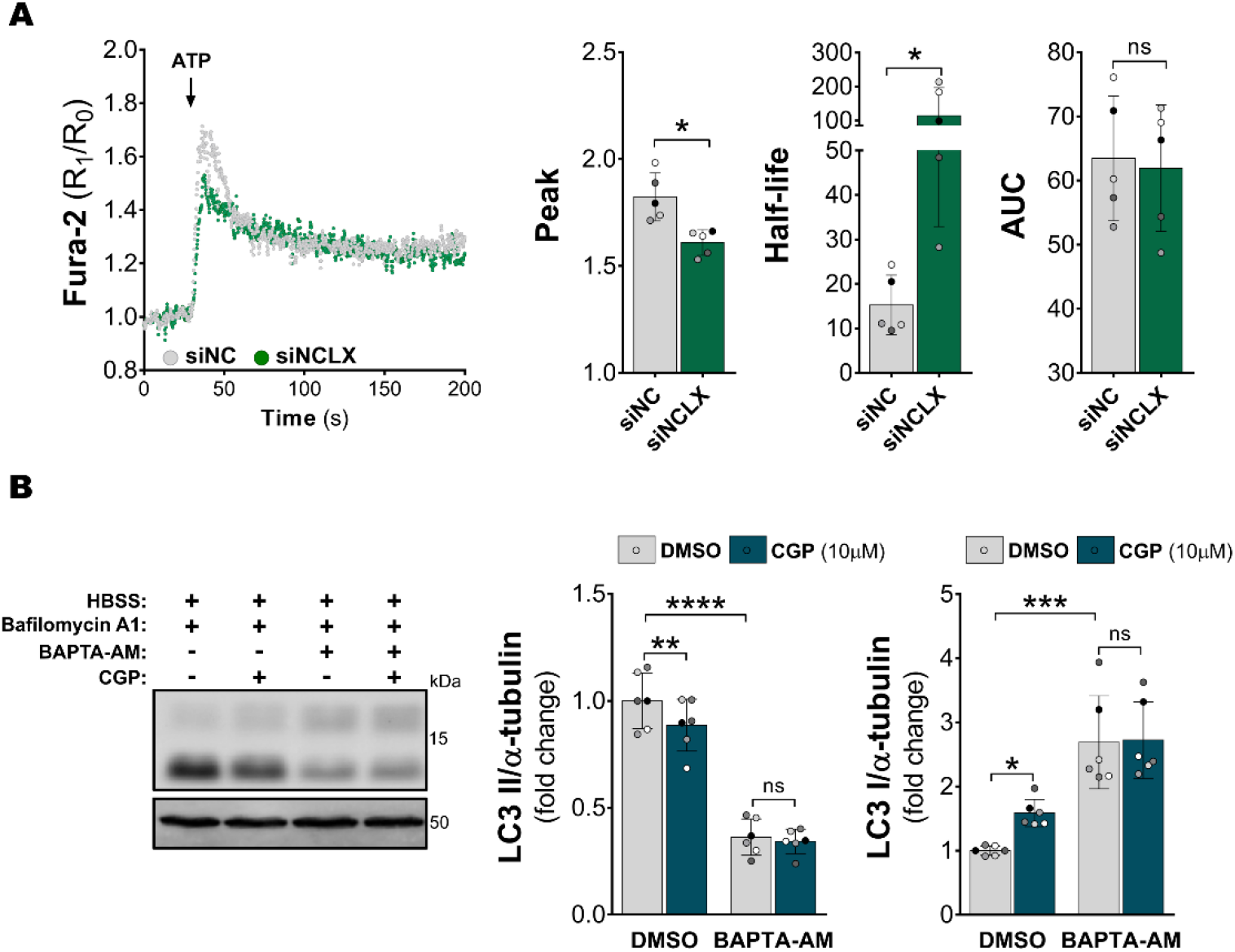
(**A**) Cytosolic Ca^2+^ measurements using Fura-2 AM after stimulation with 100 µM ATP; bar graphs showing quantifications of Ca^2+^ peaks, the half-life of one phase decay after the peak and area under the curve (AUC) (n = 5). Data were analyzed by two-tailed paired t tests. (**B**) Western blots of LC3 I and LC3 II in AML12 cells pre-incubated with 50 µM BAPTA-AM for 1 hour prior to incubation with HBSS for 1 hour in the presence either DMSO or CGP (n = 6). Data were analyzed by two-way RM ANOVA followed by Tukey’s multiple comparisons test. Data are presented in columns with means ± SD. * = p < 0.05, ** = p < 0.01, *** = p < 0.001, ns = not significant.

**Supplementary Table 1.**
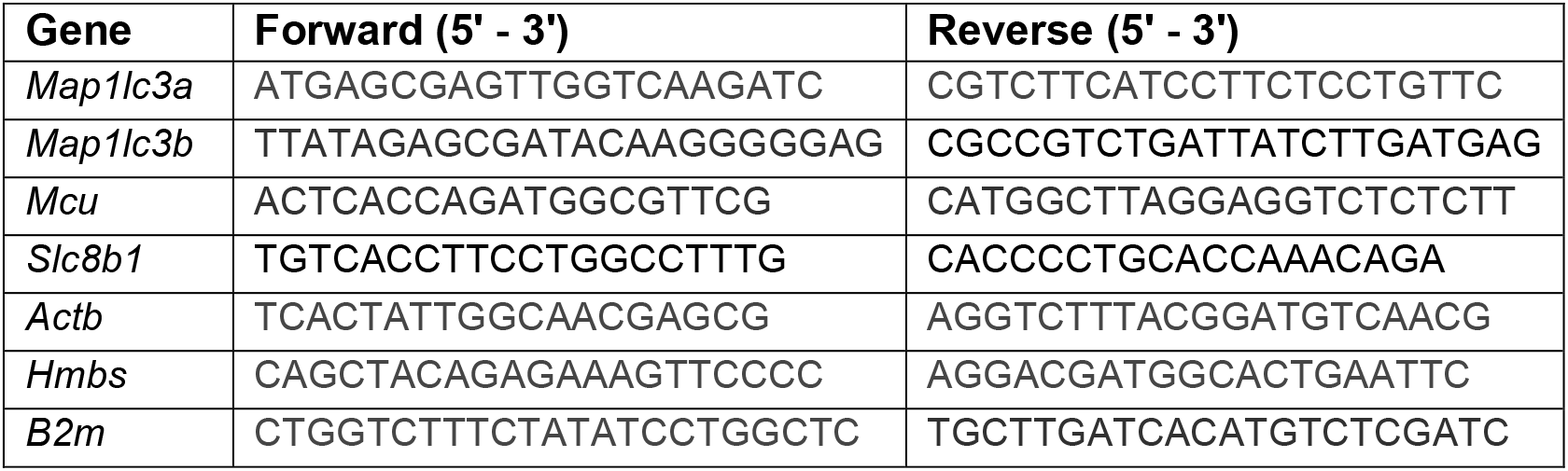
RT-qPCR primers.

